# Measuring neurofilament light in human plasma and cerebrospinal fluid: a comparison of five analytical immunoassays

**DOI:** 10.1101/2025.05.05.652212

**Authors:** Udit Sheth, Rebecca Harrison, Kyle Ferber, Erin G. Rosenbaugh, Amanda Bevis, Rohini Khillan, Michael Benatar, Nicole L. Bjorklund, Elena Di Daniel, Glenn A. Harris, Olga I. Kahn, Yongge Liu, Henrik Zetterberg, Laura L. Mitic, Danielle Graham, Tania F. Gendron

**Affiliations:** Department of Neuroscience, Mayo Clinic, 4500 San Pablo Road, Jacksonville, FL, 32224, United States; Mayo Clinic Graduate School of Biomedical Sciences, Mayo Clinic, 4500 San Pablo Road, Jacksonville, FL, 32224, United States; Astex Pharmaceuticals, 436 Cambridge Science Park Milton Rd, Milton, Cambridge, CB4 0QA, United Kingdom; Biogen, Department of Biostatistics, 225 Binney Street, Cambridge, MA, 02142, United States; Foundation at the National Institute of Health Inc., 11400 Rockville Pike, Suite 600, North Bethesda, MD, 20852, United States; Department of Neurology and ALS Center, University of Miami Miller School of Medicine, Miami, FL, 33136, United States; Association for Frontotemporal Degeneration, 2700 Horizon Drive, Suite 120, King of Prussia, PA, 19406, United States; Rainwater Charitable Foundation, 777 Main Street, Suite 2250, Fort Worth, TX, 76102, United States; Alector, Inc., 131 Oyster Point Blvd, #600, South San Francisco, CA, 94080, United States; Otsuka Pharmaceutical Development & Commercialization, Inc, 2440 Research Blvd, Rockville, MD, 20850, United States; Department of Psychiatry and Neurochemistry, Institute of Neuroscience and Physiology, the Sahlgrenska Academy at the University of Gothenburg, Mölndal, Sweden; Clinical Neurochemistry Laboratory, Sahlgrenska University Hospital, Mölndal, Sweden; Department of Neurodegenerative Disease, UCL Institute of Neurology, Queen Square, London, United Kingdom; UK Dementia Research Institute at UCL, London, United Kingdom; Hong Kong Center for Neurodegenerative Diseases, InnoHK, Hong Kong, China; Wisconsin Alzheimer’s Disease Research Center, University of Wisconsin School of Medicine and Public Health, University of Wisconsin-Madison, Madison, WI, United States; The Bluefield Project to Cure Frontotemporal Dementia and the University of California, San Francisco, San Francisco, CA 94158, United States; Biogen, Department of Biomarkers and Systems Biology, 225 Binney Street, Cambridge, MA, 02142, United States

**Keywords:** amyotrophic lateral sclerosis, biomarker, cerebrospinal fluid, immunoassays, neurofilament light chain, plasma

## Abstract

**Objectives:** Neurofilament light (NfL) is an established biofluid marker of neuroaxonal injury for neurological diseases. Several high-throughput and sensitive immunoassays have been developed to quantify NfL in blood and cerebrospinal fluid (CSF), facilitating the use of NfL as a biomarker in research and clinical practice. However, because of the lack of rigorous comparisons of assays, it has been difficult to determine whether data are comparable and whether assay performance differs. Here, we compared the performance of five NfL immunoassays.

**Methods:** To assess the five NfL immunoassays (Fujirebio, ProteinSimple, Quanterix, Roche and Siemens), we used pooled plasma or pooled CSF, as well as unique samples from 20 healthy controls and 20 individuals with El Escorial defined probable or definite amyotrophic lateral sclerosis (ALS), to evaluate precision, parallelism and/or bias. We also examined correlations between plasma and CSF NfL concentrations within and across assays and evaluated their ability to differentiate healthy controls from individuals with ALS.

**Results:** Four of the five assays demonstrated exemplary performance based on our analyses of precision and parallelism. Across the five assays, NfL concentrations were lower in plasma than in CSF, although they displayed a high degree of correlation. We noted bias across assays; plasma NfL concentrations were lowest for the Roche assay and highest for the ProteinSimple assay. In addition, all assays reliably distinguished healthy controls from individuals with ALS using plasma or CSF NfL.

**Conclusions:** Four NfL assays demonstrated similar analytic performance. Alongside performance, other factors such as costs, accessibility, useability, footprint, and intended use, should be considered.

## Introduction

Neurofilament proteins, which include neurofilament light (NfL), neurofilament medium and neurofilament heavy, are among the most widely characterized fluid biomarkers across neurodegenerative diseases. Expressed only in neurons, these type IV intermediate filaments are major components of the cytoskeletal structure and are especially abundant in myelinated axons (1). Since neurofilament proteins are released from their canonical intracellular environment in the brain into the interstitial fluid during neuroaxonal injury, their concentrations can be measured in cerebrospinal fluid (CSF) and blood. Because of the relatively easy means of measuring neurofilament proteins, they have emerged as a useful biomarker of neuronal injury and neurodegeneration. This is particularly true for NfL and its cleavage products because their greater stability and abundance maximize measurement sensitivity (2, 3). With the development of high-sensitivity analytical instruments that reliably quantify NfL in blood, which is more practical than CSF in human subjects, studies examining plasma or serum NfL across numerous neurological disorders have risen significantly over the past few years.

The greatest strength of NfL as a biomarker is its versatility. For instance, blood NfL has demonstrated prognostic utility for amyotrophic lateral sclerosis (ALS) (4), coronavirus disease 2019 (5), frontotemporal dementia (FTD) (6), Parkinson’s disease (7), stroke (8), multiple sclerosis (MS) (9) and many other neurological diseases (3, 10); nonetheless, it must be acknowledged that the prognostic potential of NfL can vary widely among neurodegenerative diseases. NfL concentrations in biofluids are also increasingly being used as a response biomarker in clinical trials (3). One such example is the VALOR study (NCT02623699), in which the efficacy and safety of tofersen, a superoxide dismutase 1 (*SOD1)*-targeting antisense oligonucleotide, was tested in clinically manifest patients with *SOD1*-ALS (11). Based on the results of this trial and an open-label extension study showing reductions in plasma NfL along with SOD1 protein reductions and improvement in clinical outcomes (11), the United States Food and Drug Administration provided accelerated approval of tofersen for the treatment of *SOD1*-ALS. Another clinical trial, the ATLAS study (<u>NCT04856982</u>), is evaluating whether tofersen delays the phenoconversion to clinically manifest ALS in pre-symptomatic carriers with highly penetrant *SOD1* variants associated with rapidly progressive disease (12). Pivotal to the ATLAS study is the use of plasma NfL as a biomarker for predicting this phenoconversion, with a predefined increase in NfL concentration triggering participant enrollment into the randomized, double-blind, and placebo-controlled period of the study. In addition to the reported utility of NfL as a prognostic, response and susceptibility/risk biomarker, it is also likely to show value in other ways. For example, it may be used as a safety biomarker in clinical trials to monitor and predict toxicity of the drug being tested (13, 14).

Given the importance of NfL for neurodegenerative disease research and the emerging potential relevance to clinical practice, we systematically compared the performance of five immunoassays for the measurement of NfL in plasma and CSF.

## Methods

### Immunoassays

Five immunoassays and corresponding instruments used to measure NfL concentrations were included in this study: (1) the Lumipulse *G* NfL blood and CSF assays run on the Fujirebio Diagnostics Inc. LUMIPULSE® G1200; (2) the Simple Plex Human NF-L Cartridge run on the ProteinSimple Ella^TM^ Automated ELISA; (3) the NF-Light v2 Advantage assay run on the Quanterix Simoa® HD-X analyzer; (4) the Elecsys® NfL run on the Roche Diagnostics cobas® e801; and (5) the Atellica NfL assay run on the Siemens Healthcare Diagnostics Atellica® IM analyzer. For simplicity, the instruments/assays are henceforth respectively referred to as Fujirebio, ProteinSimple, Quanterix, Roche, and Siemens.

### Sample procurement

To compare assay performance, we purchased matching deidentified plasma and CSF samples from 20 participants with El Escorial defined probable or definite ALS and from 20 healthy controls from PrecisionMed’s inventory of samples collected under IRB-approved clinical protocols. Plasma was derived from deidentified participant blood collected in K_2_ ethylene diamine tetra-acetic acid (EDTA) tubes, which were gently inverted ten to fifteen times, then cooled in an ice bath prior to centrifugation at 1,200 *g* at four degrees Celsius for ten minutes. The resulting plasma samples were frozen within two to three hours. Participant CSF was collected by lumbar puncture. After discarding the first one to two milliliters, CSF was collected by gravity into low protein-binding polypropylene tubes using an atraumatic needle, then centrifuged at 1,200 *g* at room temperature for ten minutes immediately after collection. The resulting blinded plasma and CSF aliquots were shipped on dry ice by PrecisionMed to Fujirebio Europe N.V., Quanterix Corporation and Siemens Healthcare Diagnostics for measuring NfL in biofluids using their respective assay, to Labcorp where NfL was measured using the Roche assay, and to Charles River Laboratories, where NfL was measured using the ProteinSimple assay.

In addition to plasma and CSF samples from controls or individuals with ALS, the University of Gothenburg created eight samples using pooled plasma from deidentified patients with high endogenous NfL to evaluate parallelism; the same was done for CSF. Blinded aliquots of these pooled samples were shipped on dry ice to PrecisionMed, who distributed them to the companies performing the NfL measurements.

### Statistical analyses

Plasma and CSF NfL concentration data provided by the above-mentioned companies were analyzed in a blinded manner. Unblinding occurred only after all statistical analyses were complete. Analyses were carried out independently by two people and concordance in the results for each assay was confirmed.

#### Evaluating repeatability and intermediate precision of plasma NfL measures

Companies used assay-specific protocols, calibrators and quality controls to report measurements for NfL in replicate using plasma and CSF from 40 participants (20 controls and 20 individuals with ALS). Plasma NfL measurements were performed across three different days, and all but one assay (Roche) had at least two runs per day (either performed by a different analyst or using a different instrument). All assays had at least two technical replicates within each run. Measurements of CSF NfL were carried out across one run, with two technical replicates in that run. When analyzing data from each assay, accommodations were made to account for differences in the number of replicates/runs among assays.

For each sample, repeatability and intermediate precision were estimated by fitting a random effects model using the lme4 package in R with restricted maximum likelihood (REML) to estimate intra-run variance, inter-run variance and overall mean concentration. Repeatability (intra-assay precision) assesses the precision of plasma NfL concentrations measured from the same sample during a single analytical run. Intermediate precision (inter-assay precision) considers variations between multiple measures of a single sample across runs and under different conditions (e.g., different instrument, different analysts). Both repeatability and intermediate precision are quantified using the coefficient of variation (%CV).

To evaluate repeatability, the residual standard deviation (SD) estimate was used to determine the sample %CV; see Equation 1. To evaluate intermediate precision, an estimate of total variance was used; see Equation 2. The overall repeatability and intermediate precision were calculated for all samples by reporting the mean and median of the sample-level repeatability and intermediate precision.

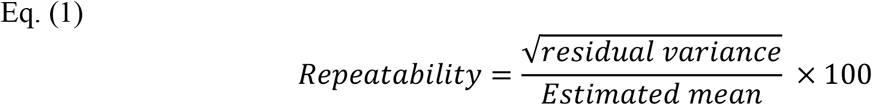

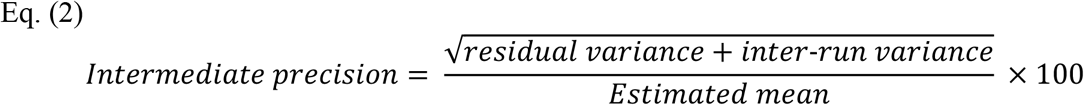

#### Evaluating plasma and CSF NfL parallelism

To evaluate parallelism, eight plasma samples and eight CSF samples with high endogenous NfL concentrations were used to measure NfL in neat samples and in samples with a range of dilutions within a given assay’s standard curve range. Note that, for ProteinSimple, comparisons were made using the 1:2 diluted samples as the reference because neat samples were depleted. Typically, plasma and CSF samples were diluted two- or four-fold, but dilutions ranged as high as 16-fold for plasma and 400-fold for CSF for one of the assays (Quanterix). The dilution series were tested in duplicate using assay-specific protocols, calibrators and quality control samples. To evaluate parallelism, NfL concentrations were corrected for the dilution factor and the %CV was calculated separately for each plasma or CSF sample. To determine the percent recovery of endogenous NfL, the sample-level percent recovery was calculated as the concentration at a given dilution divided by the mean concentration at all dilutions. The average percent recovery was calculated as the mean of the percentage recovery across all samples at a given dilution. Again, for ProteinSimple only, average percent recovery was based on the 1:2 diluted samples being the reference.

#### Correlations between plasma and CSF NfL concentrations across assays

For each assay, we evaluated Spearman’s rank correlation between mean plasma and mean CSF NfL concentrations per sample in the combined groups of controls and patients with ALS (n=40), and separately for controls (n=20) and for patients with ALS (n=20).

#### Correlations of plasma or CSF NfL concentrations across assays

We evaluated plasma NfL correlations or CSF NfL correlations across assays using samples from all 40 participants combined, and separately for each phenotype group (controls or ALS), using Spearman’s rank correlation of the mean NfL concentration per sample. A 95% confidence interval (CI) for each Spearman correlation was estimated using the nonparametric bootstrap.

#### Comparisons of bias and agreement between assays

We evaluated bias (mean difference) and limits of agreement for plasma or CSF NfL concentrations among assays using the Bland-Altman analysis. To standardize analysis methodology across assays, the average NfL measurement across replicates for each of the 40 individuals (excluding outliers) was used. Outliers were defined as a measurement of zero, a measurement below the limit of quantification, or an exceedingly high measurement orders of magnitude above the other measurements for that individual.

#### Comparisons of plasma or CSF NfL concentrations between phenotype groups across assays

For each assay, we compared plasma or CSF NfL concentrations between phenotype groups (controls and patients with ALS). Again, the average NfL measurement across replicates for each individual was used after excluding aforementioned outliers. For each assay, we tested differences in plasma or CSF NfL concentration between controls and patients with ALS using two-sided, unpaired t-tests.

In addition, to assess the ability of plasma or CSF NfL to discriminate controls from patients with ALS for each assay, we estimated area under receiver operating characteristic curve (AUC) values, where an AUC equal to 0.5 represents no discriminatory ability and an AUC equal to 1.0 represents perfect discrimination. The AUC values were calculated in R using a generalized linear model of case/control status regressed against sample mean NfL concentrations. No additional adjustments for covariates were performed.

## Results

### Immunoassay characteristics

In **Table 1**, we provide immunoassay characteristics. Each company measured NfL concentrations in the plasma and CSF samples provided to them, and we used these data to evaluate assay precision, parallelism, bias, correlations of plasma or CSF NfL concentrations between assays, correlations of plasma and CSF concentrations within each assay, and the ability of plasma or CSF NfL to distinguish patients with ALS from controls.

**Table 1:**
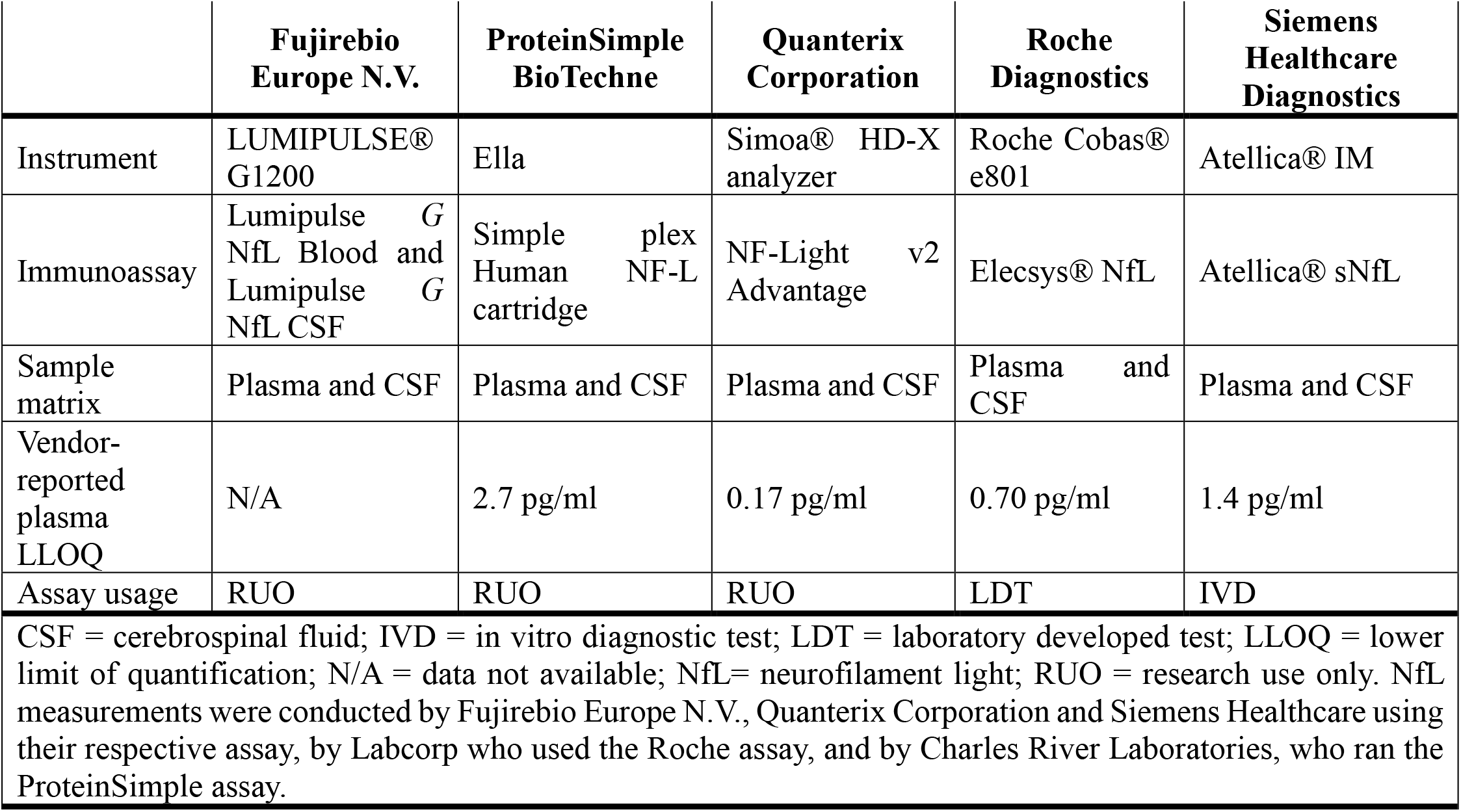
Assay Characteristics.

#### Evaluating repeatability and intermediate precision in plasma across NfL assays

To determine assay precision, we used NfL concentrations in plasma from participants with ALS and from healthy controls to evaluate repeatability and intermediate precision. For each assay, we report mean and median %CVs for repeatability and for intermediate precision in **Table 2**. Four of the five assays – Fujirebio, Quanterix, Roche and Siemens – displayed excellent repeatability with mean %CVs ranging from 2.13% to 4.77%, and median %CVs ranging from 1.60% to 4.34%. Mean and median repeatability %CVs for the ProteinSimple assay were 15.33% and 9.98%, respectively. For intermediate precision, all assays, besides ProteinSimple, reported a mean %CV≤6.94% and a median %CV≤6.11%. Mean and median intermediate precision %CVs for ProteinSimple were 20.73% and 11.46%, respectively (**Table 2**).

**Table 2:**
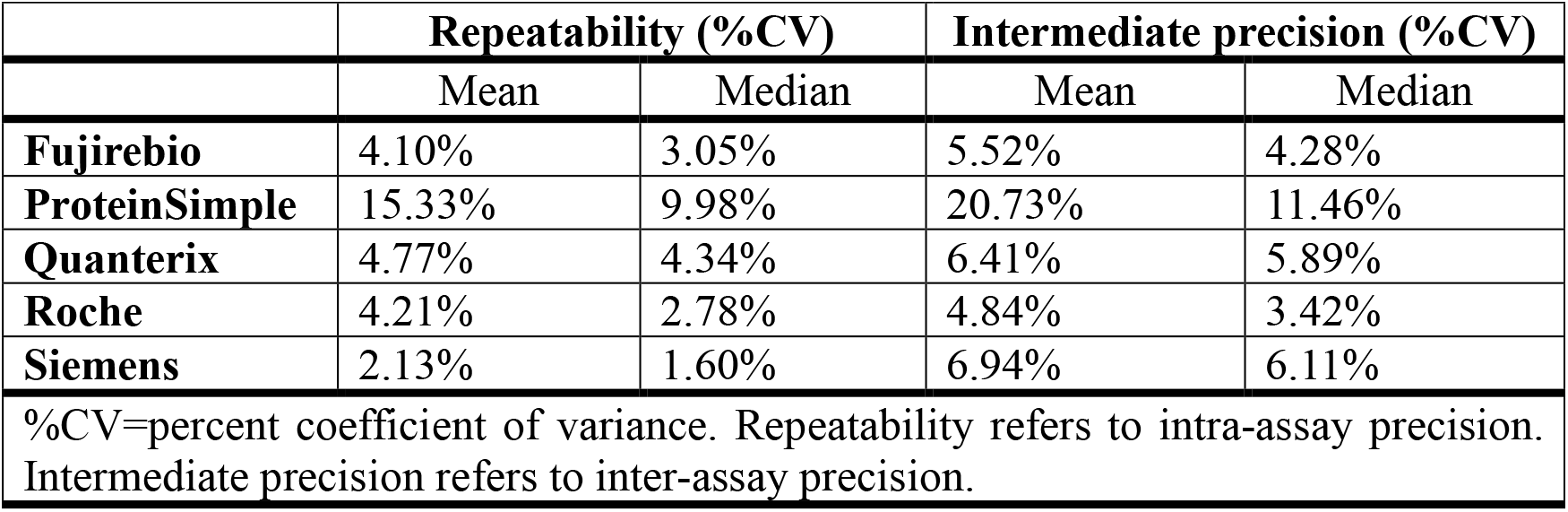
Plasma NfL repeatability and intermediate precision.

|  | **Repeatability (%CV)** |  | **Intermediate precision (%CV)** |  |
| --- | --- | --- | --- | --- |
|  | Mean | Median | Mean | Median |
| **Fujirebio** | 4.10% | 3.05% | 5.52% | 4.28% |
| **ProteinSimple** | 15.33% | 9.98% | 20.73% | 11.46% |
| **Quanterix** | 4.77% | 4.34% | 6.41% | 5.89% |
| **Roche** | 4.21% | 2.78% | 4.84% | 3.42% |
| **Siemens** | 2.13% | 1.60% | 6.94% | 6.11% |
| %CV=percent coefficient of variance. Repeatability refers to intra-assay precision. Intermediate precision refers to inter-assay precision. |  |  |  |  |

#### Evaluating plasma and CSF parallelism across NfL assays

Using pooled plasma or pooled CSF samples, we next examined parallelism, an assessment of the effect of dilution on NfL concentrations, across assays. For ProteinSimple, comparisons were made using 1:2 diluted samples as the reference. For all other assays, comparisons were made to neat samples. Optimal dilutions of the pooled plasma or CSF samples differed across assays and matrices. In **Figure 1** and **Figure S1**, we illustrate dilution factor-corrected NfL concentrations in plasma or CSF samples for each assay, and we provide mean and median %CVs for parallelism in **Table S1**. Overall, dilutions had little effect on calculated NfL concentrations. Between matrices, we observed lower %CVs in CSF than in plasma, with mean %CVs in plasma and CSF being below 10% for all assays except for ProteinSimple. The Roche assay had the lowest mean %CVs in plasma (3.93%) and in CSF (2.07%). We additionally determined the mean percent recovery of NfL concentrations in plasma or CSF samples at a given dilution; across assays, these ranged from 88% to 114% for plasma, and from 95% to 111% in CSF, with a range of 80% to 120% deemed acceptable (**Table 3**).

**Table 3:**
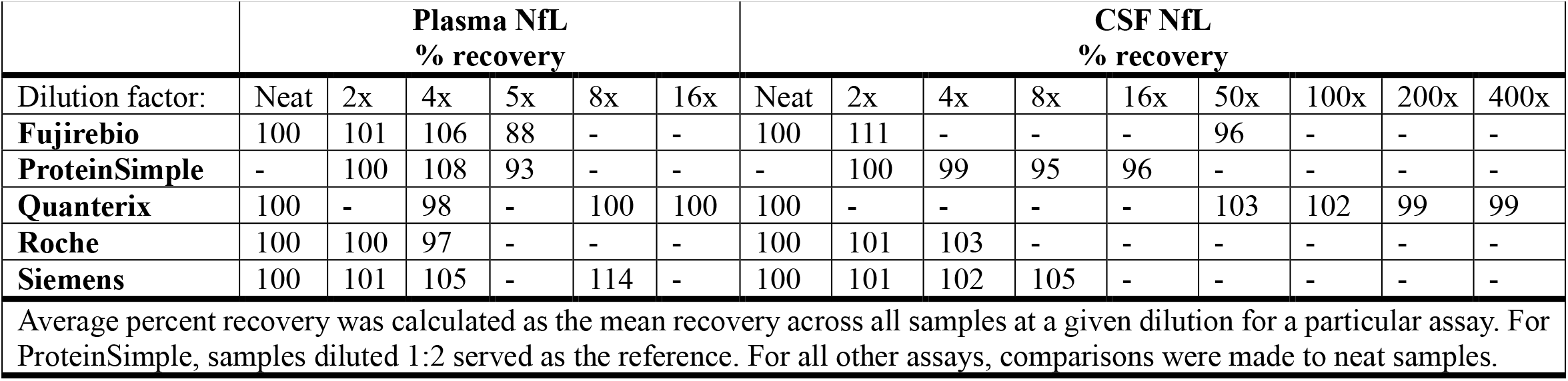
Plasma and CSF NfL recovery across diluted samples.

**Figure 1:**
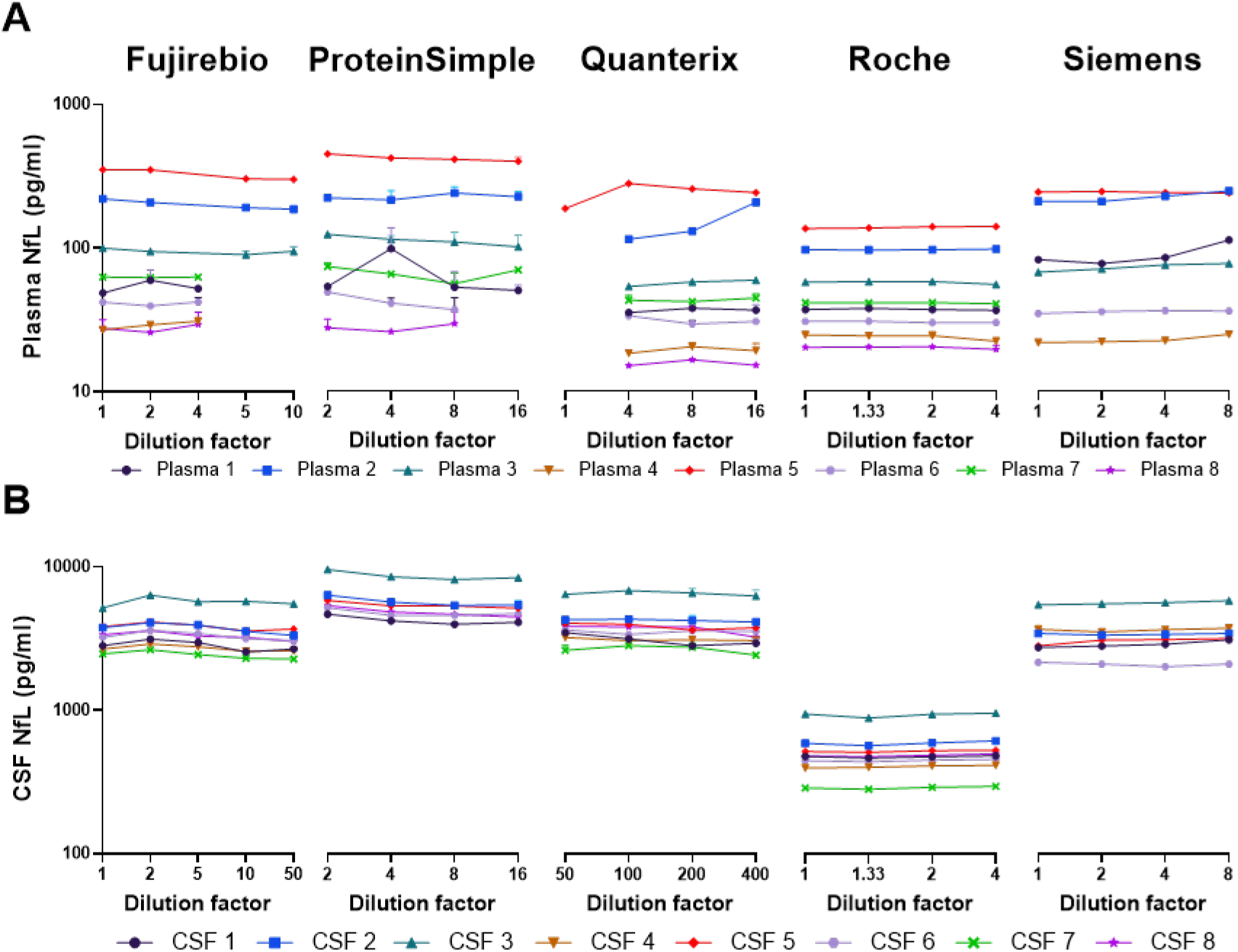
Parallelism measures for plasma and CSF NfL concentrations. NfL measurements across a series of manufacturer-recommended dilutions using pooled plasma (**A**) or CSF (**B**) samples. Data are shown on the Log10 scale for ease of visualization. For data shown on the linear scale, please refer to **Figure S1**.

#### Evaluating correlations between plasma and CSF NfL concentrations

For each assay, we evaluated correlations between plasma and CSF NfL concentrations in the combined groups of controls and patients with ALS (n=40), finding strong positive correlations (Spearman’s r: 0.91 for all assays) (**Table S2**). When comparing each group separately, Spearman’s r values were generally lower in controls (n=20), ranging from 0.66 to 0.76 across assays, and higher in patients with ALS (n=20) with Spearman’s r values ranging from 0.70 to 0.83.

#### Evaluating correlations of plasma or CSF NfL concentrations and estimating bias between assays

We examined correlations of NfL concentrations in plasma or CSF across the five assays, finding strong and highly statistically significant correlations (**Figure 2, Table 4**). Indeed, Spearman’s r values ranged from 0.96 to 0.99 in plasma, and from 0.99 to 1.00 in CSF. However, when visualizing the correlations (**Figure 2**), bias (mean difference) between assays was apparent, and this was corroborated by Bland-Altman analysis. Among all pair-wise comparisons of assays, bias between assays was notable except when comparing plasma or CSF NfL concentrations between the Quanterix and Siemens assays (**Figures S2 and S3**, and **Table S3**).

**Table 4:** Correlations of plasma or CSF NfL concentrations between assays.

| **Plasma correlations** |  | **Fujirebio** | **ProteinSimple** | **Quanterix** | **Roche** | **Siemens** |
| --- | --- | --- | --- | --- | --- | --- |
| **Fujirebio** | r<br>p-value | 1.00 |  |  |  |  |
| **ProteinSimple** | r<br>p-value | 0.98<br><2.2E-16 | 1.00 |  |  |  |
| **Quanterix** | r<br>p-value | 0.99<br><2.2E-16 | 0.98<br><2.2E-16 | 1.00 |  |  |
| **Roche** | r<br>p-value | 0.99<br><2.2E-16 | 0.98<br><2.2E-16 | 0.99<br><2.2E-16 | 1.00 |  |
| **Siemens** | r<br>p-value | 0.98<br><2.2E-16 | 0.96<br><2.2E-16 | 0.97<br><2.2E-16 | 0.98<br><2.2E-16 | 1.00 |
| **CSF correlations** |  | **Fujirebio** | **ProteinSimple** | **Quanterix** | **Roche** | **Siemens** |
| **Fujirebio** | r<br>p-value | 1.00 |  |  |  |  |
| **ProteinSimple** | r<br>p-value | 1.00<br><2.2E-16 | 1.00 |  |  |  |
| **Quanterix** | r<br>p-value | 0.99<br><2.2E-16 | 0.99<br><2.2E-16 | 1.00 |  |  |
| **Roche** | r<br>p-value | 1.00<br><2.2E-16 | 0.99<br><2.2E-16 | 0.99<br><2.2E-16 | 1.00 |  |
| **Siemens** | r<br>p-value | 1.00<br><2.2E-16 | 0.99<br><2.2E-16 | 0.99<br><2.2E-16 | 1.00<br><2.2E-16 | 1.00 |

r=Spearman's correlation coefficient

**Figure 2:**
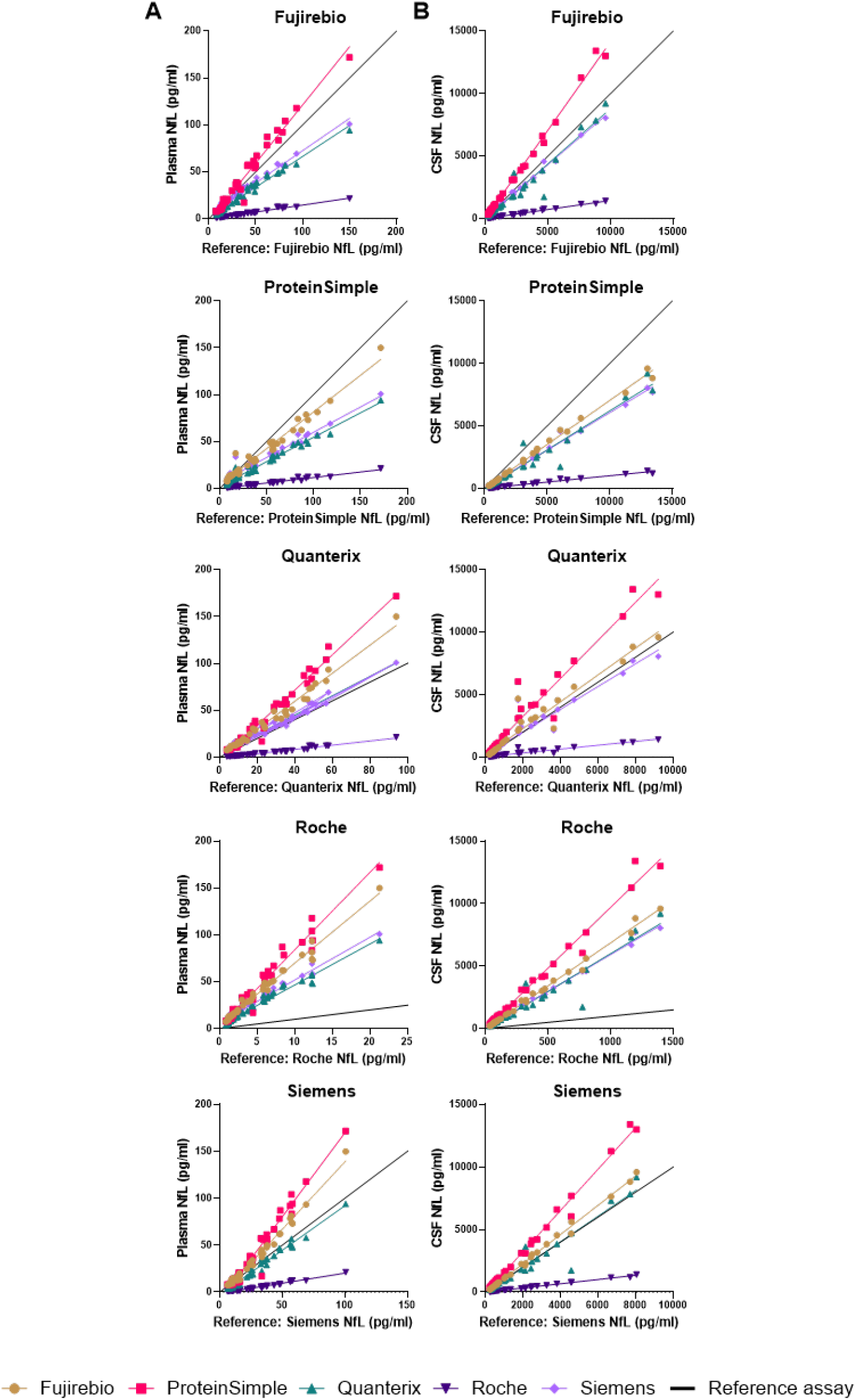
Correlations of plasma or CSF NfL concentrations between assays. Correlations of plasma NfL (**A**) or CSF NfL (**B**) concentrations across all five assays. Assay names at the top of each graph serve as the reference assay, depicted by a black line, to which the other assays are compared. Fujirebio=gold, ProteinSimple=pink, Quanterix=green, Roche=purple, Siemens=lavender, reference assay=black.

#### Evaluating plasma or CSF NfL concentrations between phenotype groups

For each assay, we compared plasma or CSF NfL concentrations between controls and patients with ALS finding statistically significant increases in NfL in patients with ALS compared to controls (p<0.0001) for all assays (**Figure 3, Figure S4**). Finally, we observed that both plasma and CSF NfL discriminated controls from individuals with ALS with high AUC values ranging from 0.88 to 0.95 for plasma NfL, and from 0.92 to 0.99 for CSF NfL. Plasma and CSF NfL AUC values were highest for the Fujirebio assay (**Table 5**).

**Table 5:** Plasma and CSF NfL performance in discriminating controls from individuals with ALS.

|  | **Plasma NfL** |  |  | **CSF NfL** |  |  |
| --- | --- | --- | --- | --- | --- | --- |
| **Assay** | AUC | 95% CI | p-value | AUC | 95% CI | p-value |
| **Fujirebio** | 0.95 | 0.91-0.99 | <0.0001 | 0.99 | 0.96-1.00 | <0.0001 |
| **ProteinSimple** | 0.89 | 0.76-1.00 | <0.0001 | 0.93 | 0.84-1.00 | <0.0001 |
| **Quanterix** | 0.89 | 0.77-1.00 | <0.0001 | 0.92 | 0.83-1.00 | <0.0001 |
| **Roche** | 0.88 | 0.76-1.00 | <0.0001 | 0.93 | 0.84-1.00 | <0.0001 |
| **Siemens** | 0.88 | 0.75-0.99 | <0.0001 | 0.93 | 0.84-1.00 | <0.0001 |
| AUC= area under the receiver operating curve; CI=confidence interval |  |  |  |  |  |  |

**Figure 3:**
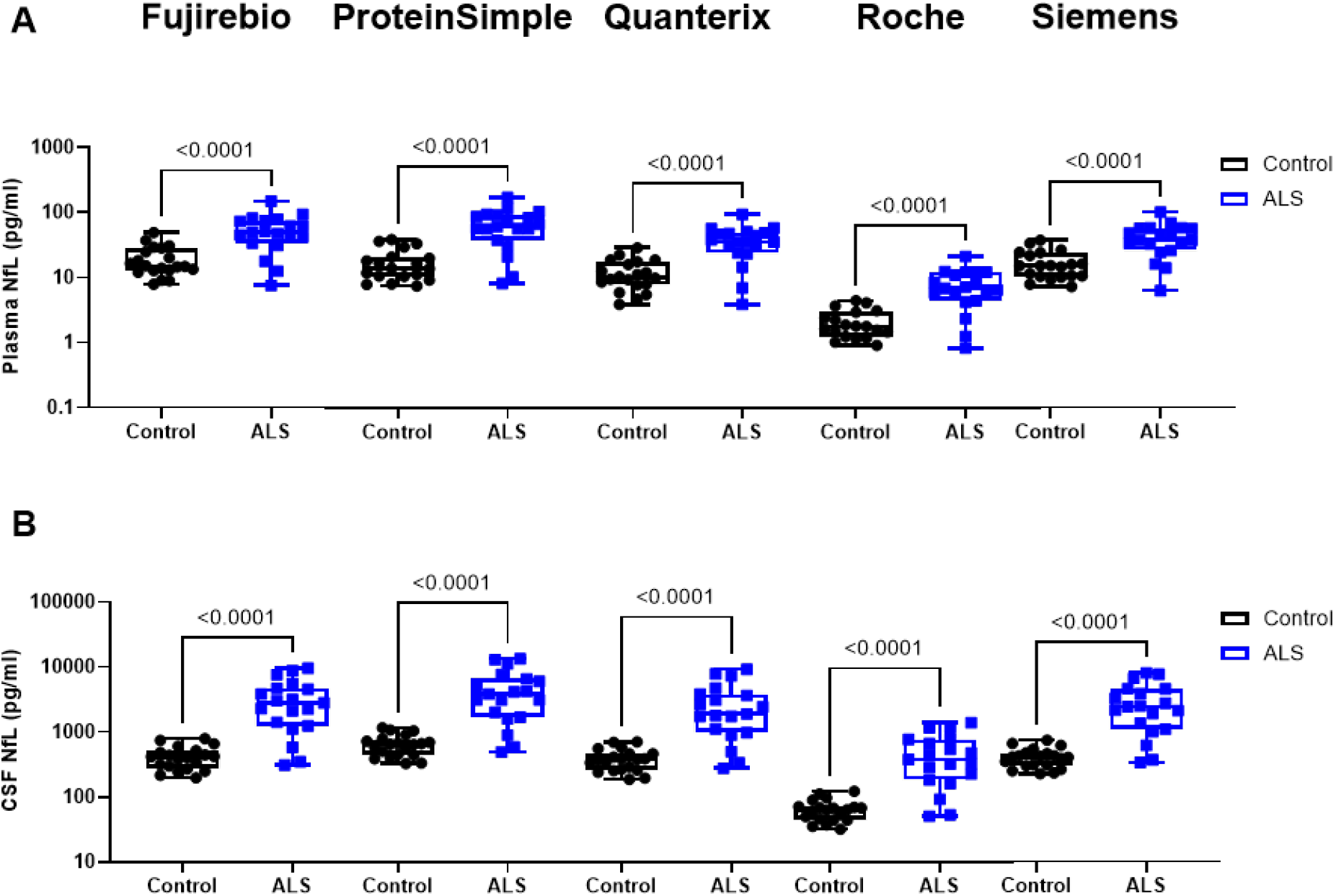
Plasma and CSF NfL concentrations in controls and patients with ALS. For each assay, plasma NfL (**A**) and CSF NfL (**B**) concentrations in 20 controls and 20 patients with ALS were compared. Maximum and minimum NfL concentrations are represented by the top and bottom whiskers, respectively. The 75% and 25% quartiles are represented by the upper and lower bounds of the box, respectively, and the median is represented by the mid-line of the box. Data are shown using a Log10 scale for ease of visualization. P-values from two-sided, unpaired t-tests are shown. For data visualized on the linear scale, please refer to **Figure S4**.

## Discussion

It is well recognized that CSF and blood NfL are markers of neuroaxonal damage for a range of neurological disorders. Because this biomarker boasts various utilities, it is increasingly used in research and in some clinical settings, especially now that ultrasensitive assays for the quantification of blood NfL have been developed (15). Here, we compared NfL precision and parallelism across assays. In addition to these analytical characteristics, we examined assay bias, NfL concentration correlations, and the ability of NfL to distinguish healthy controls from patients with ALS.

Regarding repeatability (intra-assay precision) and intermediate precision (inter-assay precision) in plasma, all assays performed similarly except for ProteinSimple, which showed less consistency. Whereas ProteinSimple mean repeatability and intermediate precision %CVs were above 15%, the other assays demonstrated mean repeatability and intermediate precision %CVs lower than 7%. Our data are largely congruent with those from a recent study in which intra-assay and inter-assay %CVs were less than 8% when measuring NfL in serum using the Fujirebio, ProteinSimple, Quanterix and Roche assays (16).

We additionally observed that mean %CVs for parallelism were highest for ProteinSimple being greater than 15% for plasma but below 10% for CSF. Among the other four assays, mean %CVs for parallelism were below 10% for both plasma and CSF. Across all assays, mean percent NfL concentration recovery ranged from 88% to 114% in plasma, and from 95% to 111% in CSF.

For each of the five assays, plasma and CSF NfL concentrations correlated strongly in the combined group of controls and patients with ALS. However, when evaluating correlations of plasma and CSF NfL for each phenotype group separately, Spearman’s r correlation coefficients were generally lower in controls than in patients with ALS.

Correlations of plasma or CSF NfL concentrations between all five assays were very strong. However, we did note a bias among assays; plasma NfL concentrations were lowest for the Roche assay and highest for the ProteinSimple and Fujirebio assays. These findings are in line with results from a prior study that performed a head-to-head comparison of plasma NfL concentrations using Fujirebio, Quanterix, Roche and Siemens assays (17), and the aforementioned study in which serum NfL was quantified using the Fujirebio, ProteinSimple, Quanterix and Roche assays (16), and underscore that absolute values for NfL quantification are not readily comparable across assays. Differences in NfL concentrations among assays could be due to differences in the antibody pairs being used for each assay.

When comparing plasma or CSF NfL concentrations between controls and patients with ALS, we observed that plasma and CSF NfL were statistically significantly increased in patients with ALS across all assays. We also observed that both plasma and CSF NfL discriminated controls from individuals with ALS with high AUC values (plasma: 0.88-0.95; CSF: 0.92-0.99). Importantly, however, we do not interpret these results to support the utility of NfL as a diagnostic tool since the case-control design we employed, comparing equal number of patients and controls, does not reflect the diagnostic challenge encountered in clinical practice.

The strengths of our study include the blinding of samples used for NfL quantification, the blinding of instruments/assays when analyzing data, the inclusion of healthy control samples, and the comparison of five NfL assays. While our study was ongoing, two similar studies were published; one in which NfL assays from Fujirebio, Quanterix, Roche and Siemens were compared (17). In that study, the authors assessed plasma NfL correlations and bias between assays and compared the distribution of plasma NfL concentrations between assays using three disease groups: patients initially diagnosed with ALS, patients with ALS having undergone treatment, and patients with MS. In a similar fashion, the second study compared NfL assays from Fujirebio, ProteinSimple, Quanterix, and Roche, and assessed serum NfL concentration correlations among assays and compared serum NfL concentrations between patients with or without ALS (16). In the present study, we assessed both plasma and CSF NfL concentration correlations and bias among assays and evaluated plasma and CSF NfL between phenotype groups. Moreover, we used two parameters to measure precision across assays and two parameters to measure parallelism, providing a more robust comparison of performance across assays. We also evaluated the ability of plasma or CSF NfL concentrations to distinguish controls from patients with ALS for each assay.

Our study also has some limitations. When analyzing precision for each assay, the number of replicates or runs among assays were not uniform and, when analyzing parallelism, some assays (ProteinSimple and Siemens) tested only six of the eight samples for a given matrix. Finally, for ProteinSimple, samples diluted 1:2 served as the reference for parallelism studies in contrast to using neat samples for all other assays.

In summary, our data demonstrate that the Fujirebio, Quanterix, Roche and Siemens assays showed greater precision than the ProteinSimple assay. However, all immunoassays similarly distinguished controls from patients with ALS, and for all other comparisons, the five assays performed relatively similarly. The high correlations of NfL concentrations across assays bode well for currently ongoing standardization studies that will be instrumental when using NfL in clinical settings (18). It must also be noted that, while we present objective data, additional factors must too be considered when choosing the ideal NfL immunoassay; these include the investigators’ intended use, instrument and reagent costs, required sample volume, instrument size, degree of automation, and whether the assay is designated for research use only or as an *in vitro* diagnostic or a laboratory developed test.

## Author contributions

All authors contributed to the study design, analysis plan, data review, manuscript review, and approval of the final manuscript for submission. KF, RH and US also contributed to data analysis. TFG and US wrote the manuscript with input from all authors.

### Acknowledgements

We thank Frank Shewmaker, Christine Swanson-Fischer and all other FNIH Biomarker Consortium members for their participation in this study.

## Research funding

The following academic or private-sector partners provided financial or in-kind support for this study: Foundation at the National Institute of Health Inc., Robert Packard Center for ALS Research at Johns Hopkins, Alector, Inc., Biogen MA, Inc., Diagnostics Accelerator at the Alzheimer’s Drug Discovery Foundation, Otsuka Pharmaceutical Development & Commercialization, Inc., Rainwater Charitable Foundation, The ALS Association, The Association for Frontotemporal Degeneration, and The Bluefield Project to Cure Frontotemporal Dementia. MB is supported by U54NS092091, U01NS107027, and R01NS10547. HZ is a Wallenberg Scholar and a Distinguished Professor at the Swedish Research Council supported by grants from the Swedish Research Council (#2023-00356, #2022-01018 and #2019-02397), the European Union’s Horizon Europe Research and Innovation Programme under grant agreement No 101053962, Swedish State Support for Clinical Research (#ALFGBG-71320), and the European Partnership on Metrology, co-financed from the European Union’s Horizon Europe Research and Innovation Programme and by the Participating States (NEuroBioStand, #22HLT07).

## Data availability

De-identified NfL biomarker data are available upon request.

## Consent of sample usage

To compare assay performance, we purchased matching de-identified plasma and CSF samples from 20 participants with El Escorial defined probable or definite ALS and from 20 healthy controls from PrecisionMed’s inventory of samples collected under IRB-approved clinical protocols and eight pooled plasma and eight pooled CSF samples from University of Gothenburg collected under IRB-approved protocols.

## List of abbreviations

ALS: amyotrophic lateral sclerosis
AUC: area under the receiver operating curve
CI: confidence interval
CSF: cerebrospinal fluid
%CV: percent coefficient of variance
EDTA: ethylene diamine tetra-acetic acid
FTD: frontotemporal dementia
MS: multiple sclerosis
NfL: neurofilament light
r: Spearman’s correlation coefficient
REML: restricted maximum likelihood model
SD: standard deviation
SOD1: superoxide dismutase 1

