## Supplemental Figures and Tables for "Measuring neurofilament light in human plasma and cerebrospinal fluid: a comparison of five analytical immunoassays"

**Supplementary figures and tables**

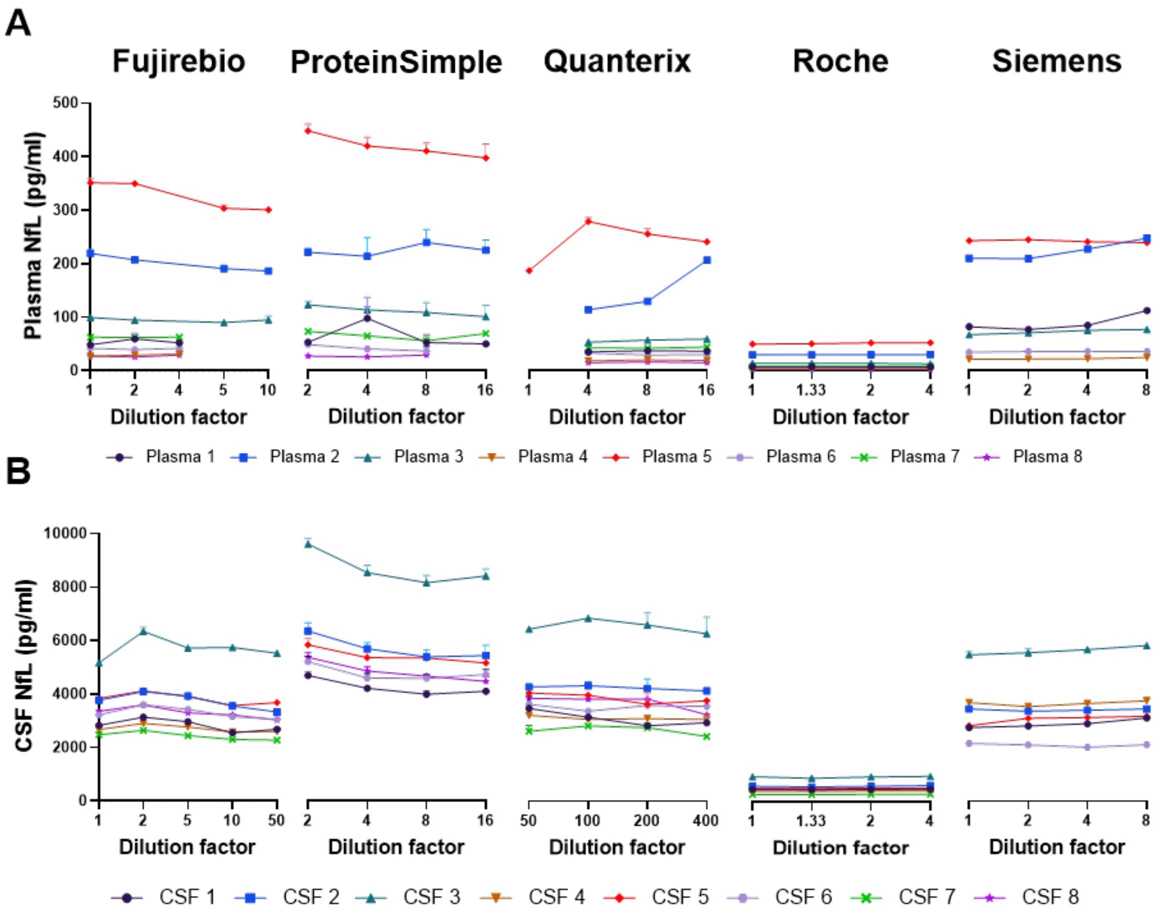


**Figure S1: Parallelism measures for plasma and CSF NfL concentrations.** NfL measurements across a series of manufacturer-recommended dilutions using pooled plasma (**A**) or CSF (**B**) samples. Data are shown on the linear scale.

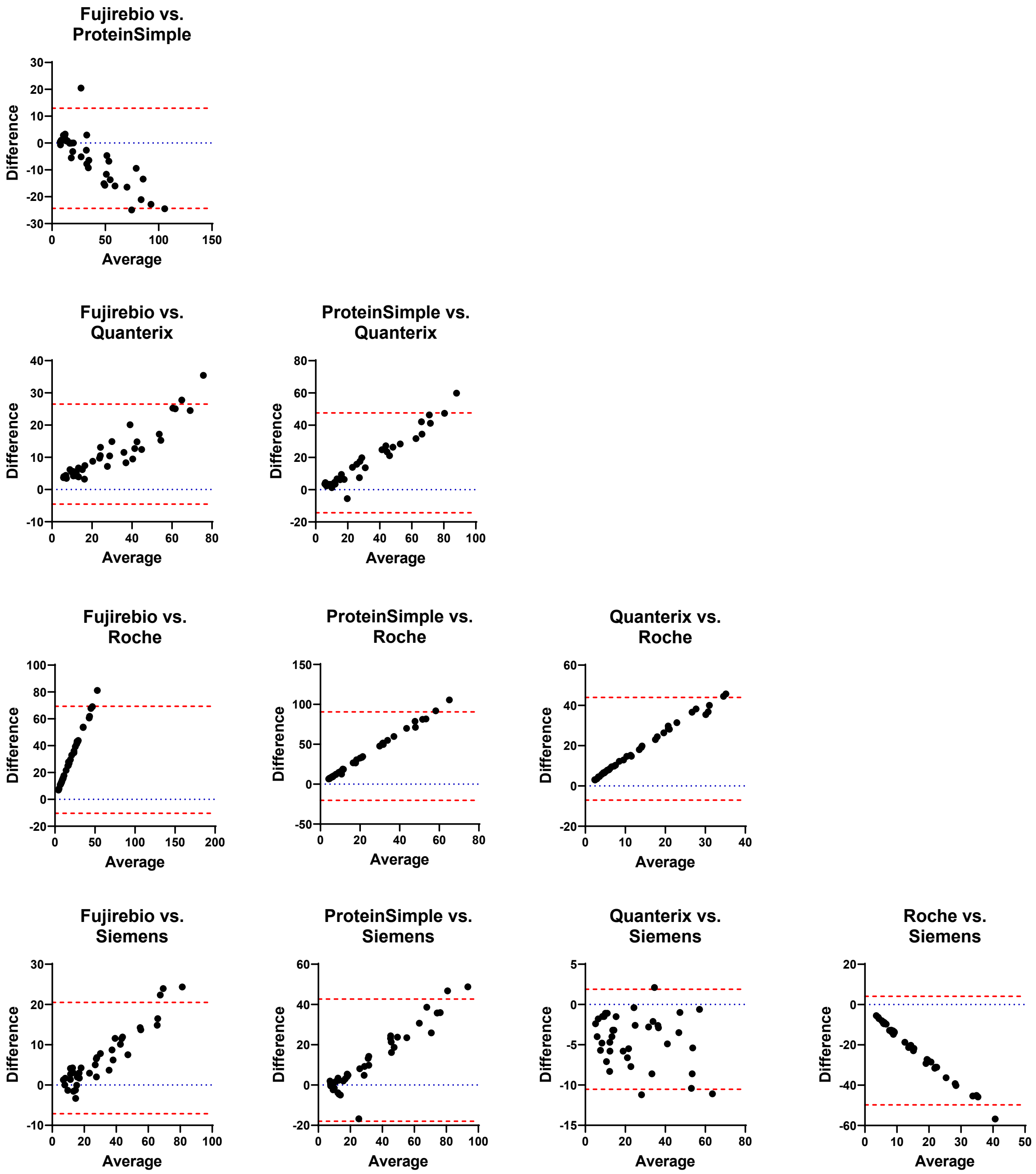


**Figure S2: Pair-wise comparisons of assays measuring plasma NfL.** Bland-Altman plots showing the comparison of two assays used to measure NfL in plasma samples. The blue dotted line represents Y=0 (the difference between the two NfL concentrations being compared), while the red dotted lines indicate the 95% limits of agreement.

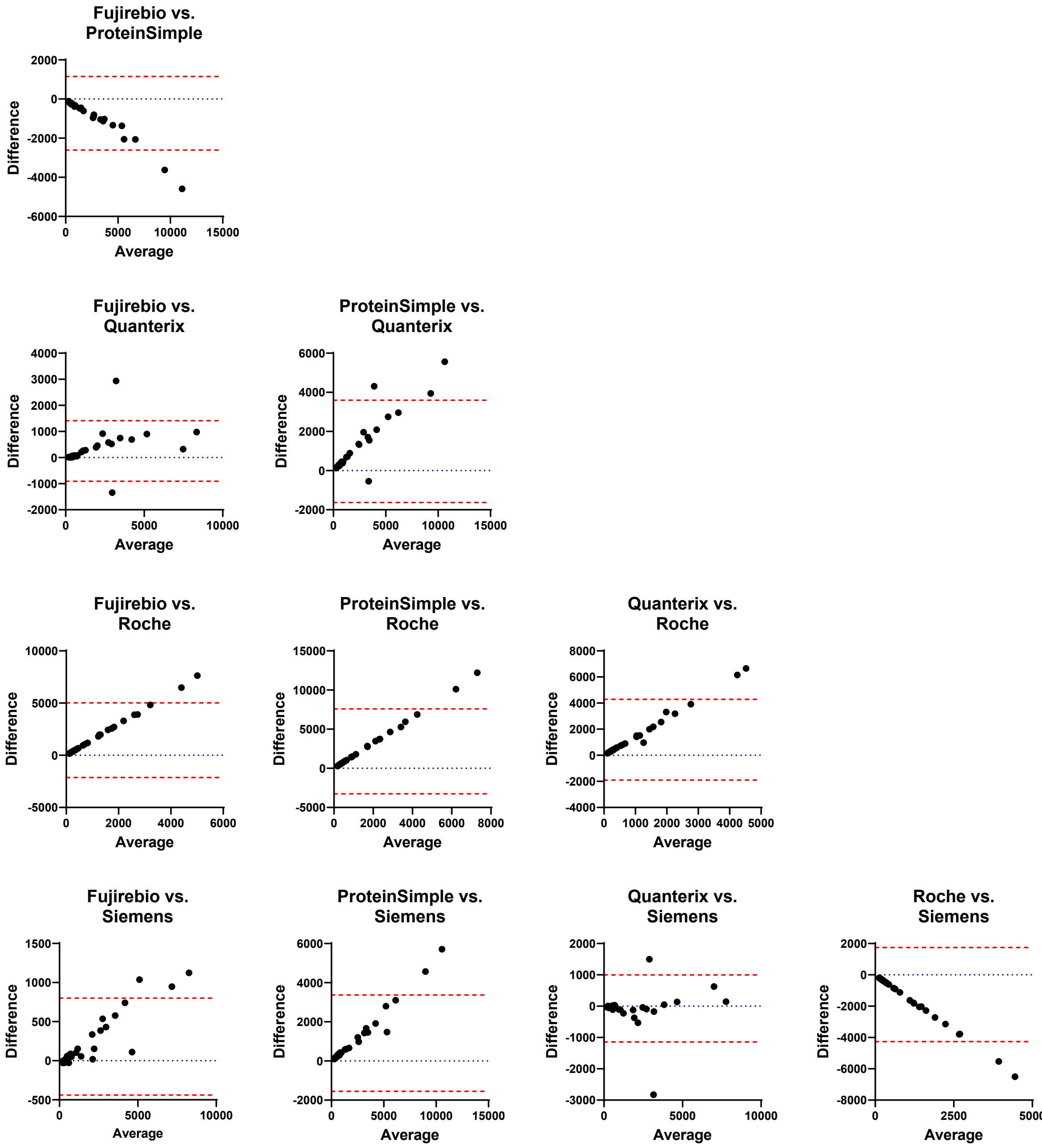


**Figure S3: Pair-wise comparisons of assays measuring CSF NfL.** Bland-Altman plots showing the comparison of two assays used to measure NfL in CSF samples. The blue dotted line represents Y=0 (the difference between the two NfL concentrations being compared), while the red dotted lines indicate the 95% limits of agreement.

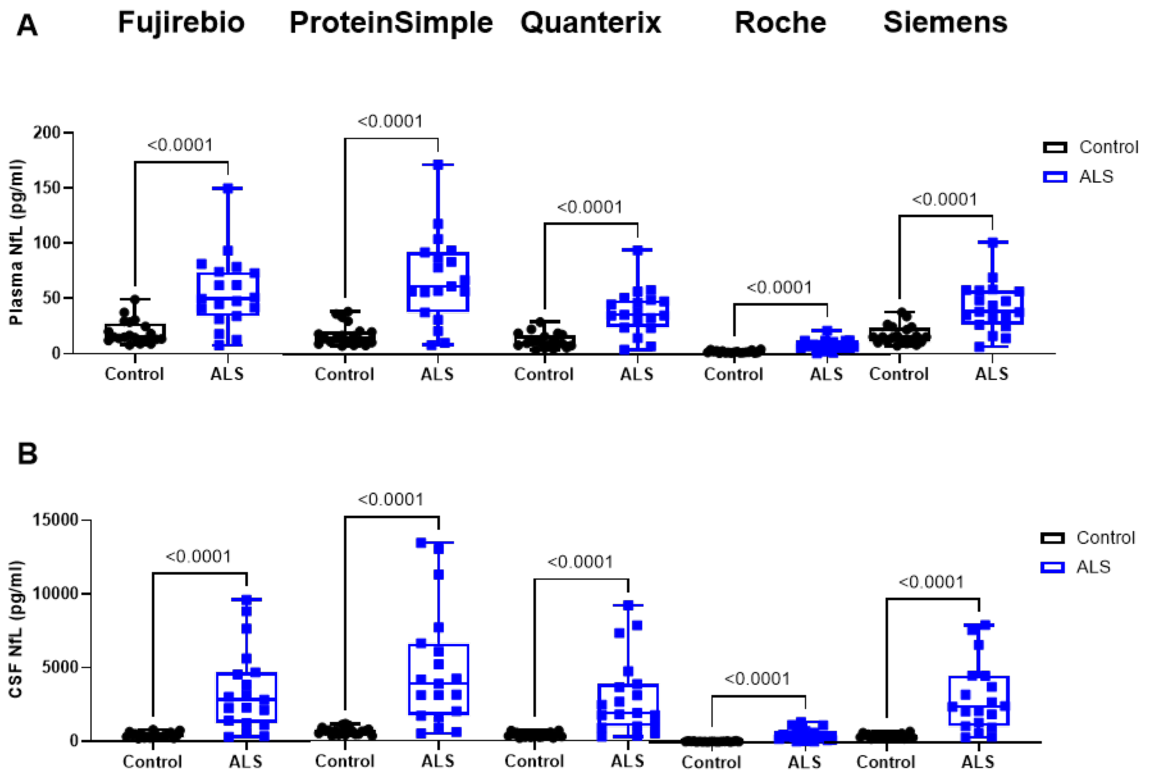


**Figure S4: Plasma and CSF NfL concentrations in controls and patients with ALS.** For each assay, plasma NfL (**A**) and CSF NfL (**B**) concentrations in 20 controls and 20 patients with ALS were compared. Maximum and minimum NfL concentrations are represented by the top and bottom whiskers, respectively. The 75% and 25% quartiles are represented by the upper and lower bounds of the box, respectively, and the median is represented by the mid-line of the box. Data are shown on the linear scale. P-values from two-sided, unpaired t-tests are shown.

**Table S1: Mean and median parallelism %CVs across assays**

| | **Plasma parallelism (%CV)** | | **CSF parallelism (%CV)** | |
|---|---|---|---|---|
| | Mean | Median | Mean | Median |
| **Fujirebio** | 7.40% | 6.82% | 6.43% | 6.34% |
| **ProteinSimple** | 17.53% | 13.11% | 6.99% | 6.97% |
| **Quanterix** | 9.65% | 6.02% | 5.89% | 5.86% |
| **Roche** | 3.93% | 3.24% | 2.07% | 1.86% |
| **Siemens** | 6.60% | 5.84% | 3.34% | 2.83% |

CV=coefficient of variance. For ProteinSimple, samples diluted 1:2 served as the reference. For all other assays, comparisons were made to neat samples.

**Table S2: Spearman correlations between plasma and CSF NfL concentrations across phenotype groups and assays**

| | **All participants combined** | | **Controls** | | **ALS** | |
|---|---|---|---|---|---|---|
| | r | p-value | r | p-value | r | p-value |
| **Fujirebio** | 0.91 | <0.0001 | 0.73 | 0.0003 | 0.80 | <0.0001 |
| **ProteinSimple** | 0.91 | <0.0001 | 0.66 | 0.0016 | 0.83 | <0.0001 |
| **Quanterix** | 0.91 | <0.0001 | 0.73 | 0.0002 | 0.70 | 0.0006 |
| **Roche** | 0.91 | <0.0001 | 0.68 | 0.0009 | 0.76 | <0.0001 |
| **Siemens** | 0.91 | <0.0001 | 0.76 | <0.0001 | 0.78 | <0.0001 |

r=Spearman correlation coefficient. Calculated for plasma and CSF NfL measures within a given assay for both phenotype groups combined or for each group separately.

**Table S3: Bland-Altman analysis of plasma or CSF NfL among assays**

| Plasma bias | | Fujirebio | ProteinSimple | Quanterix | Roche | Siemens |
|---|---|---|---|---|---|---|
| **Fujirebio** | Bias<br>95% LoA | | | | | |
| **ProteinSimple** | Bias<br>95% LoA | -5.67<br>(-24.33, 12.99) | | | | |
| **Quanterix** | Bias<br>95% LoA | 11.00<br>(-4.54, 26.54) | 16.67<br>(-14.26, 47.60) | | | |
| **Roche** | Bias<br>95% LoA | 29.49<br>(-10.36, 69.34) | 35.16<br>(-20.24, 90.56) | 18.49<br>(-7.01, 43.98) | | |
| **Siemens** | Bias<br>95% LoA | 6.69<br>(-7.13, 20.51) | 12.36<br>(-17.99, 42.70) | -4.31<br>(-10.52, 1.90) | -22.80<br>(-49.74, 4.13) | |
| **CSF bias** | | **Fujirebio** | **ProteinSimple** | **Quanterix** | **Roche** | **Siemens** |
| **Fujirebio** | Bias<br>95% LoA | | | | | |
| **ProteinSimple** | Bias<br>95% LoA | -729.2<br>(-2609, 1150) | | | | |
| **Quanterix** | Bias<br>95% LoA | 253.6<br>(-90.62, 1413) | 982.8<br>(-1630, 3595) | | | |
| **Roche** | Bias<br>95% LoA | 1443<br>(-2132, 5017) | 2172<br>(-3253, 7597) | 1189<br>(-1907, 4285) | | |
| **Siemens** | Bias<br>95% LoA | 181.4<br>(-438.4, 801.2) | 910.6<br>(-1549, 3370) | -72.18<br>(-1144, 999.6) | -1261<br>(-4265, 1743) | |

Bias=average difference between sample reads between two given assays. LoA = limits of agreement. Bias and LoA represent column-row values.
